# Hue selectivity from recurrent circuitry in *Drosophila*

**DOI:** 10.1101/2023.07.12.548573

**Authors:** Matthias P Christenson, Álvaro Sanz Díez, Sarah L Heath, Maia Saavedra-Weisenhaus, Atsuko Adachi, L.F. Abbott, Rudy Behnia

## Abstract

A universal principle of sensory perception is the progressive transformation of sensory information from broad non-specific signals to stimulus-selective signals that form the basis of perception. To perceive color, our brains must transform the wavelengths of light reflected off objects into the derived quantities of brightness, saturation and hue. Neurons responding selectively to hue have been reported in primate cortex, but it is unknown how their narrow tuning in color space is produced by upstream circuit mechanisms. To enable circuit level analysis of color perception, we here report the discovery of neurons in the *Drosophila* optic lobe with hue selective properties. Using the connectivity graph of the fly brain, we construct a connectomics-constrained circuit model that accounts for this hue selectivity. Unexpectedly, our model predicts that recurrent connections in the circuit are critical for hue selectivity. Experiments using genetic manipulations to perturb recurrence in adult flies confirms this prediction. Our findings reveal the circuit basis for hue selectivity in color vision.

## Introduction

Perceived features can differ significantly from the physical attributes of sensory stimuli. These differences play a crucial role in how animals interact with the world around them. Understanding the neural circuit basis of the transformation from physical detection to perception is central to neuroscience. In color vision, the relationship between the spectral composition of light and derived color percepts is particularly dramatic. Reflectance spectra of objects in our environment are high dimensional, varying in infinite combinations of wavelengths and intensity. In contrast, our perception of color is low dimensional, varying only along the three axes of hue, saturation and brightness. Hue, saturation and brightness are perceptual variables, that each are derived from, but are not equal to, specific aspects of the physical properties of light. The hue or tint of a color is related to the mean wavelength composition of the spectrum. Saturation or degree of purity of a color is related to the variance of the spectrum. Brightness is related to the total intensity of the spectrum. How these stimulus-selective perceptual variables are computed in the brain and emerge from activation of photoreceptors in the eyes remains to be determined.

In primate color vision, cone photoreceptor excitations combine into color opponent signals that ultimately are transformed into hue selective responses (Thompson et al., 1992; Mehrani et al., 2020; Flachot and Gegenfurtner, 2021). Hue selective neurons throughout visual cortex are narrowly tuned to specific spectral hues such as cyan, teal and orange, or non-spectral hues such as purple and magenta, and are relatively independent of other features such as saturation and brightness (Hanazawa et al., 2000; Horwitz and Hass, 2012; Conway et al., 2007; Lennie et al., 1990; Komatsu et al., 1992; Zeki, 1980). Other neurons show selectivity for particular saturation or brightness levels (Hanazawa et al., 2000; Xing et al., 2015; Li et al., 2022). While primates are well suited to define the role of these particular signals in perception, many challenges remain when working with such complex brains to unravel the circuit mechanisms that lead to the construction of these signals. Recently however, the opportunity to study this question in circuits that are genetically accessible and mapped at the EM connectome level has arisen in the brain of the fruit fly *Drosophila melanogaster* (Schnaitmann et al., 2018; Zimmermann et al., 2018; Heath et al., 2020).

In the fruit fly *Drosophila melanogaster*, color vision begins in yellow and pale variants of R7 and R8 photoreceptors (pR7, yR7, pR8, yR8) that express one of four opsins (Rh3, Rh4, Rh5 and Rh6, respectively) with peak sensitivities in the short UV, UV, blue or green. As in trichromatic primates, fly photoreceptor activations are combined into color opponent signals that already emerge at the terminals of the R7 and R8 photoreceptors (Schnaitmann et al., 2018; Heath et al., 2020). Color opponency is the result of axo-axonal interactions between photoreceptors and additional connections mediated by Dm9 interneurons. We now ask how these opponent signals are further transformed downstream of the photoreceptors. Although the main postsynaptic partners of R7 and R8 have been identified (Li et al., 2021; Kind et al., 2021) and some have been implicated in color guided behaviors (Melnattur et al., 2014; Karuppudurai et al., 2014), the chromatic response properties of these neurons have not been described previously. Here we take advantage of the genetic tractability that fruit flies afford to measure visual responses of candidate neurons across fly color space. We show that three transmedulary (Tm) projection neurons downstream of R7s and R8s, Tm5a, Tm5b and Tm20, have tunings that differ from those of their photoreceptor inputs, with maximal responses to violet and various non-spectral colors. Furthermore, the response of Tm5a, Tm5b and Tm20 show selectivity to hue.

To leveraged this discovery and explore the circuit mechanisms behind these Tm responses, we analyzed fly connectome data (Zheng et al., 2018; Dorkenwald et al., 2023; Schlegel et al., 2023) and constructed a circuit-level model based on this analysis and on our measurements of additional interneuron responses. This model accurately matches the Tm responses we measured. In addition, the model allowed us to predict the effects of silencing different neurons in the circuit, predictions that we verified through genetic silencing experiments. We find that recurrent connections within the circuit are critical for the color tuning and hue selectivity of Tm neurons. This work identifies biological mechanisms that govern the transition from sensory detection to perceptual representation in the brain.

## Results

### Hue selectivity is defined by the geometry of responses in color space

We begin by discussing human trichromatic color vision, both because it is more familiar and because its color space can be depicted more easily in figures than the fly’s tetrachromatic system. The three types of cone photoreceptors in humans, S, M and L (Fig.1a), define the three axes shown in Figure 1b. Any visible color can be mapped to a point in this three-dimensional space located at distances along each axis equal to the activation of the corresponding photoreceptor. For colors with the same luminance, the sum of these activations is constant, and this constraint defines the chromatic triangle shown in Figure 1c, where only hue and saturation can vary. Spectral colors fall along a line within the color triangle corresponding to activation of the photoreceptors by a single wavelength of light. Equal activity across photoreceptors, corresponding to the point in the center of this triangle, is perceived as white. Any non-white color including non-spectral colors that cannot be recreated by a single wavelength of light can be represented by a vector stretching from the white point to the point corresponding to the particular color. We define hue as the angle θ of this vector relative to a given axis and saturation s as its length.

**Figure 1.**
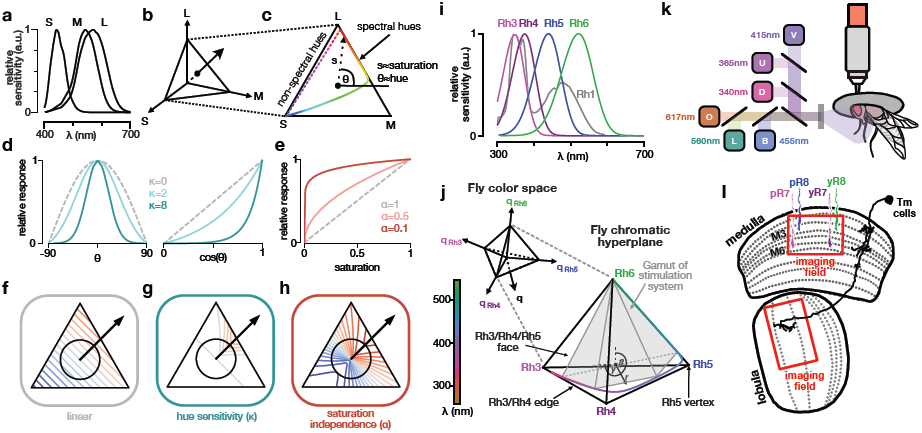
Geometry of hue selective responses in color space. **(a)** Sensitivities of the human S, M, and L cone photoreceptors (adapted from Stockman et al. (1993)). **(b)** Human trichromatic color space defined by the excitations of the S, M, and L cone. **(c)** Trichromatic chromatic hyperplane, where the vertices indicate exclusive excitation of the S, M, or L cone. The center of the triangle is the white point, the angle *θ* defines the hue of a color, and the distance from the white point *s* defines its saturation. **(d)** Different sensitivities to the hue angle relative to the preferred direction of tuning for fixed saturation. The gray dashed line shows the linear case and the colored lines show increasing sensitivity to hue. **(e)** Different sensitivity to saturation for a fixed hue angle. The gray dashed line shows the linear case and the colored lines show decreasing sensitivity to saturation. **(f)** Isoresponse curves in the trichromatic hyperplane for a linear neuron. **(g)** Isoresponse curves in the trichromatic hyperplane for a hue sensitivity neuron (*κ* = 8). **(h)** Isoresponse curves in the trichromatic hyperplane for a saturation independent neuron (*α*=0.1). **(i)** Relative spectral sensitivity of opsins expressed in the fruit fly retina; data from Salcedo et al. (1999) fit with equation from Govardovskii et al. (2000). **(j)** Fly color space defined by the captures of Rh3, Rh4, Rh5, and Rh6 opsins. The luminance of a color is defined as the sum of opsin captures. Restricting the sum of the captures to a constant value, we obtain a chromatic hyperplane. In the chromatic hyperplane, there are a total of 4 vertices, one for each opsin, six edges between pairs of opsins, and a four faces connecting three opsins. The gray box indicate the gamut of fly colors accurately reproducible with our stimulation system. Within the chromatic plane, the “saturation” of a color is equal to the distance of the stimulus from the fly white point (i.e. the center of the tetrahedron). Two angles *β* and *γ* define the “hue” of a color. **(k)** Two-photon imaging set up. The fly is secured facing LED setup, and LED sources are combined using a custom color mixer to form a single collimated beam. LED abbreviations: D: dUV; U: UV; V: violet; B: blue; L: lime; O: orange. **(l)** Schematic of the fruit fly color circuit indicating the imaging fields used to record photoreceptors and interneurons in the medulla and Tm neurons in the lobula.

The preferred color tuning of a neuron is defined as the hue direction that produces the maximum response for a given saturation. Color selectivity is then assessed by studying how the neuron’s response diminishes for colors rotated away from this preferred direction by an angle θ. For a neuron with a linear response, this reduction is proportional to cos(θ) (Fig.1d grey dashed lines). The response of such a linear neuron will also be linear across saturation s (Fig.1e grey dashed lines). As we will show, such a linear model provides a good description of the responses at the axon terminals of the photoreceptors, but not of the Tm neurons. We characterize hue selective cells by measuring how their responses deviate from the linear model. In particular, hue selective neurons are characterized by having higher sensitivity (tighter tuning) to hue angle θ (Fig.1d colored lines) and reduced sensitivity to saturation s (Fig.1e colored lines), as well as reduced sensitivity to brightness, i.e. luminance (a variable not included in the chromatic triangle). In the following, we show that these features are found in Tm5a, Tm5b and Tm20 neural responses.

To get a sense of the geometry in color space implied by hue and saturation selectivity, it is useful to consider the geometry of the corresponding response patterns in color space as a whole (Horwitz and Hass, 2012; Golden et al., 2016). A linear response can be visualized as evenly spaced parallel iso-response lines that are orthogonal to a preferred tuning direction (Fig.1f). Along a circle of constant saturation, this tuning has the cosine shape discussed above, peaking at the preferred direction of tuning (Fig.1d). In this two-dimensional framework, increased sensitivity to hue (Fig.1d) corresponds to bending iso-response contours into U shaped curves (Fig.1g), without changing the spacing between them. Reduced sensitivity to saturation (Fig.1e) produces iso-response lines that are rotated toward the preferred direction of tuning and spaced further apart, also forming U shaped curves (Fig.1h). Both of these non-linear mechanisms produce responses that are more confined in color space and contribute to hue selectivity.

The fly eye has five opsins (Fig.1i). Here we only consider the four Rh3-6 expressed in R7 and R8, as Rh1, expressed in R1-6, is broadband and implicated in achromatic vision. Thus the analog of the triangle shown in Figure 1c is, instead, a tetrahedron (Fig.1j). This means that there are two hue angles for the fly, corresponding, for example, to the polar and azimuthal angles of a three-dimensional spherical coordinate system. The preferred color direction for a given cell and the color direction for a particular stimulus are thus each characterized by two angles. Nevertheless, the difference between these directions defines a single angle θ as in the trichromatic case, and we will use this angle to define color selectivity as in Fig.1d.

Although each fly photoreceptor only expresses a single opsin, their axonal responses reflect activation from multiple opsins due to direct (axo-axonal) and indirect (through the horizontal cell Dm9) interactions that give rise to color opponency (Heath et al., 2020; Schnaitmann et al., 2018). We therefore need to distinguish between the effects of light on the opsins and on the activity at axon terminals, which is where all the photoreceptor recordings reported here were made. For circuit modeling, the axonal responses are of obvious relevance because they represent the input to downstream targets. Nevertheless, to align with convention, we use computed photon capture (relative to the background) of the opsins, rather than axonal responses, to determine the locations of points within the color tetrahedron. As a result, we label the axes of the tetrahedron Rh3, Rh4, Rh5 and Rh6.

Identifying different mechanisms of hue selectivity from their response geometries requires measurement of responses over a large portion of the color space, which is challenging. In primates, approaches such as closed loop iso-response methods (Gollisch and Herz, 2012; Horwitz and Hass, 2012) have been developed to try and circumvent this problem. In *Drosophila*, we instead take advantage of genetically identified and accessible neurons to measure the responses of particular chromatic neuron types over nearly the entire fly color space by sampling across multiple single neurons and animals. Using the fly’s spectral sensitivities, we constructed a “gamut stimulus set” that combines six LEDs (Fig.1k) at different relative intensities to cover a range of calculated photon captures that span the available chromatic dimensions in flies. We restricted our analysis to stimuli that were close to a single isoluminant hyperplane (Fig.1j shaded area). In addition, to evaluate response properties across different luminance levels, we created a “contrast stimulus set” consisting of increasing intensities of single and mixtures of LED flashes. We used both stimuli to probe the response properties of neurons in the *Drosophila* chromatic circuit, using cell type specific two-photon imaging of GCAMP6f (Fig.1k,l).

### The main downstream targets of R7 and R8 photoreceptors have diverse chromatic tuning properties that differ from their direct photoreceptor input

The main downstream targets of R7 and R8 photoreceptor axons have been identified (Kind et al., 2021): yR7 axons target Tm5a, pR7 axons target Tm5b, Tm20 is downstream of either pR8 or yR8 axons in their home column and Tm5c is downstream of either of the R8s, favoring yR8s over pR8s. We imaged the responses of the terminals of all four Tm neurons in the lobula to the gamut stimulus and compared them to those of their photoreceptor inputs.

Examination of responses across the color tetrahedron (Fig.2a-d) provides an indication of the preferred chromatic tuning of each Tm neuron. Tm5a is activated only close to the Rh3 vertex with some activation along the Rh3/Rh6 edge (Fig.2a). Tm5b is activated only along the Rh4/Rh5 edge of the tetrahedron (Fig.2b). Tm5c responses are broad with a slight preference close to the Rh6 vertex (Fig.2c). Tm20 has strongest activation within the Rh4/Rh5/Rh6 face (Fig.2d).

The three dimensional nature of the fly isoluminant color space makes it difficult to visualize color tuning (Fig.2a-d). It is useful to examine responses along onedimensional paths through the three-dimensional color space. To do this, we used radial basis functions to smoothly interpolate the responses from both the gamut and contrast stimulus sets (see Methods), allowing us to plot them along any chosen line within the color space. Along the single wavelength line, we obtain a spectral tuning curve (Fig.2 For photoreceptors (Fig.S1 and i-l), the results are, overall, consistent with our previous measurements (Heath et al., 2020). In the case of Tm neurons, the curve peaks in the UV part of the spectrum for Tm5a (Fig.2f), violet for Tm5b (Fig.2g) and blue for Tm20 (Fig.2i). Tm5c is most sensitive to violet/ blue/green light but has broader tuning than the other Tm neurons (Fig.2h).

In addition to these spectral sensitivities, we examined tuning to non-spectral colors (Goldsmith, 1990; Thompson et al., 1992; Endler and Mielke, 2005; Stoddard et al., 2020). In humans, for example, purple is the result of excitation of both S and L cones. Analogous colors for fruit flies corresponds to exciting combinations Rh3-Rh6 opsins, Rh4-Rh6 opsins, or Rh3-Rh5 opsins in various ratios (lines in Fig.2j). We define a neuron as non-spectrally tuned if its peak response is greater along a non-spectral line than along the single wavelength line. We observed such non-spectral tuning in Tm5a, Tm5c and Tm20. Tm5a has its strongest response to a relative combination of 0.9 Rh3 and 0.1 Rh6 (Fig.2k). Tm5c has strong tuning along all the non-spectral lines we investigated, with peaks at 0.1 Rh3 and 0.9 Rh5, at 0.2 Rh4 and 0.8 Rh6, and at 0.3 Rh3 and 0.7 Rh6 (Fig.2m). Tm20 has its strongest response to a relative combination of 0.2 Rh4 and 0.8 Rh6 (Fig.2n). In contrast, photoreceptor axon terminals and Tm5b have a clear and unambigious spectral response (Fig.S1o-r and Fig.2l).

If Tm5a, Tm5b and Tm20 responses were driven solely by their direct photoreceptor inputs, they would appear as sign inverted version of the photoreceptor responses as a result of their inhibitory histaminergic synapses. Instead, the responses of these neurons peak at different locations in color space. Thus, additional connections in the chromatic circuit are required to account for the Tm responses.

### Tm5a, Tm5b and Tm20 carry sparse chromatic signals

In contrast to the photoreceptor terminals, Tm5a, Tm5b and Tm20 do not show a strong response to stimuli close to the “fly white” point (Fig.2a, b and d). This suggests that they may have a reduced sensitivity to luminance. To quantify sensitivity to luminance, we used linear regression to compute how well Tm responses could be explained by the luminances of the stimuli compared to locations of the stimuli within the color tetrahedron. To do this, we used both the gamut and contrast stimulus sets. We defined the ratio of the resulting *r*^2^s as a luminance invariance index (see Methods), with larger values indicating more invariance. All the photoreceptor terminals have luminance invariance indices around 0.5 (Fig.S1g). Tm neurons indices vary across types (Fig.2o), with Tm20 showing a luminance invariance index similar to that of the photoreceptor terminals, and Tm5c having a decreased luminance invariance. Tm5b and Tm5c, on the other hand, are less sensitive to luminance than the photoreceptor terminals with luminance invariance indices around 1.2 (Fig.2o).

Finally, visual inspection of Figure 2a, b and d suggests that response of Tm5a, Tm5b and, to a lesser degree, Tm20 are sparse. To quantify sparsity, we calculated the area under a normalized cumulative histogram of the Tm responses. A uniform distribution of responses by this measure would yield a sparsity index of 0.5. Values larger than 0.5 correspond to sparser response distributions in which responses are more likely to be close to zero than close to the maximum. Tm5b has the sparsest response profile, with a sparsity index of 0.85, followed closely by Tm5a and Tm20, with indices of 0.8 and 0.75 respectively (Fig.2p). Tm5c in contrast, has a profile that is similar to those of the photoreceptor axonal responses with an index around 0.6 (Fig.S1h and Fig.2p).

**Figure 2.**
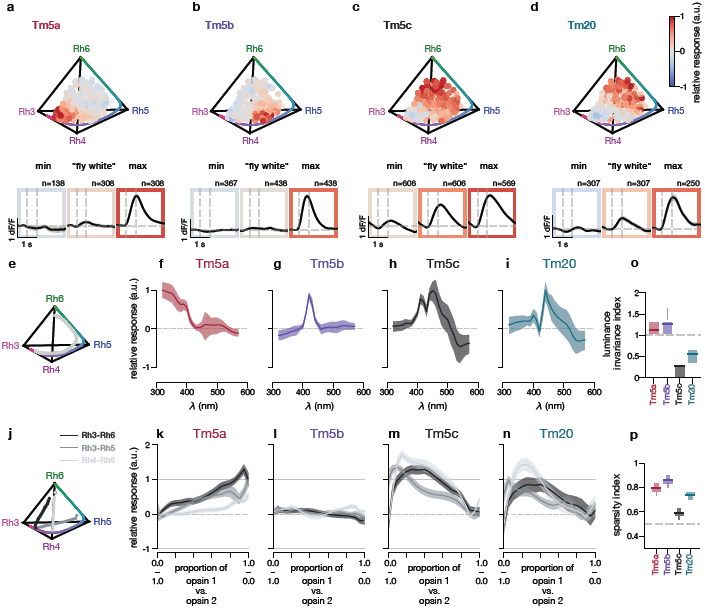
Sparse responses and non-spectral selectivity in Tm neurons. **(a-h)** Relative amplitude responses of Tm neurons across the gamut of tested fly colors. Chromatic stimuli are represented as points in the chromatic hyperplane, with the color of each point indicating the relative response of the indicated neuron to that stimulus. The colored line that spans the edges of the tetrahedron from Rh3 to Rh4, Rh5 and Rh6 corresponds to single wavelengths., ranging across all single wavelengths from 300 to 560nm (i.e. the visible spectrum of the fly). The three lower insets for each plot show dF/F responses across time from 0.5s before stimulus onset to 1.5s after stimulus offset, with onset and offset indicated by the dashed gray vertical lines. The dashed gray horizontal line indicates the baseline dF/F. From left to right, the insets show the average responses across recorded neurons around the location of the minimum amplitude, around the middle of the tetrahedron (the “fly white” point), and around the location of the maximum response. The spherical volume used for averaging responses around the desired location for each inset has a radius of 0.05, with each edge of the tetrahedron having unit length. **(e)** Approximate single wavelength line used for the interpolation of responses (line of gray scatter points). The colored line is the actual single wavelength line. **(f-i)** Interpolated single wavelengths for photoreceptors and Tm neurons. The colored line is the interpolated mean response. The shaded areas is the bootstrapped 95% percentile of the mean response. **(j)** Non-spectral lines used for interpolation of responses (lines of gray scatter points). The colored line corresponds to the pure single wavelength line. **(k-n)** Interpolated non-spectral responses of Tm neurons. The colored lines correspond to the interpolated mean responses for different non-spectral lines (as indicated in the legend). The shaded areas correspond to the bootstrapped 95% percentile of the mean response. **(o)** Sparsity indices for Tm neurons. A value of 0.5 corresponds to a uniform response distribution. The vertical histogram corresponds to the bootstrapped distribution and the line for each cell type is the mean sparsity index. **(p)** Calculated luminance invariance indices for Tm neurons. The luminance invariance index is the ratio of the goodness-of-fit (*R*^2^) of a linear regression using only the chromatic dimensions (only opponent-like interactions) and the *R*^2^ of a linear regression using only the achromatic dimension (sum of photoreceptors). The vertical histogram corresponds to the bootstrapped distribution and the line for each cell type is the mean luminance invariance index.

Our results to this point show that, downstream of photoreceptors, visual information is reformatted into four parallel channels that convey distinct types of information. Tm5a and Tm5b have non-spectral and spectral preferred tunings, respectively, have sparse responses with reduced sensitivity to luminance. Tm20 shares these features but has more sensitivity to luminance. In contrast, Tm5c is broadly tuned in color space and has a heightened sensitivity to luminance. In addition to having a response tuning that is different from the other Tms, the connectomic analysis discussed below indicates that, despite its projection to the lobula, it is equivalent to an interneuron, acting mostly locally within this circuit. For these reasons, we omit Tm5c from our analysis of hue selectivity, but we include it as a source of recurrent connections in the model that we discuss later in the paper.

### Tm5a, Tm5b and Tm20 are hue selective

The responses of Tm5a, Tm5b and, to a somewhat lesser extent, Tm20 have properties that are reminiscent of so-called “hue-selective” neurons that have been identified in the cortex of trichromatic primates (Hanazawa et al., 2000; Du et al., 2022). In the remaining sections, we focus on establishing this quantitatively, and then we explore the circuit mechanisms underlying the signal transformation that leads to hue selectivity in these Tm neurons.

In contrast to their photoreceptor inputs (Heath et al., 2020) and to Tm5c (Fig.S2a), Tm5a, Tm5b and Tm20 are not well fit by a linear model (Fig.3a). Including Rh1, which is known to provide some input to color pathways (Pagni et al., 2021) does not improve these fits, except for a slight increase in the *R*^2^ for Tm20 (Fig.S2b). Including an output non-linearity in the form of a tanh function does not improve any of the fits, with or without Rh1 (Fig.3a and Fig.S2b). This implies that some form of nonlinear processing of photoreceptor excitation occurs to give rise to the responses at the level of Tm neurons. We therefore constructed a non-linear model that, as a function of its parameters, allowed for both sharpening of sensitivity to hue angle and decreased sensitivity to saturation. The model response depends on the angle between the stimulus color vector and the neuron’s preferred color direction, *θ*, and on the saturation level *s*. In this model, to allow for enhanced hue sensitivity and reduced sensitivity to saturation, two parameter *κ* and *α* can vary respectively. The values *κ→* 0 and *α* = 1 result in a linear response, while increasing *κ* tightens the model’s color tuning, and decreasing α reduces its dependence on saturation. The curves shown in (Fig.1dh) were generated by this model.

**Figure 3.**
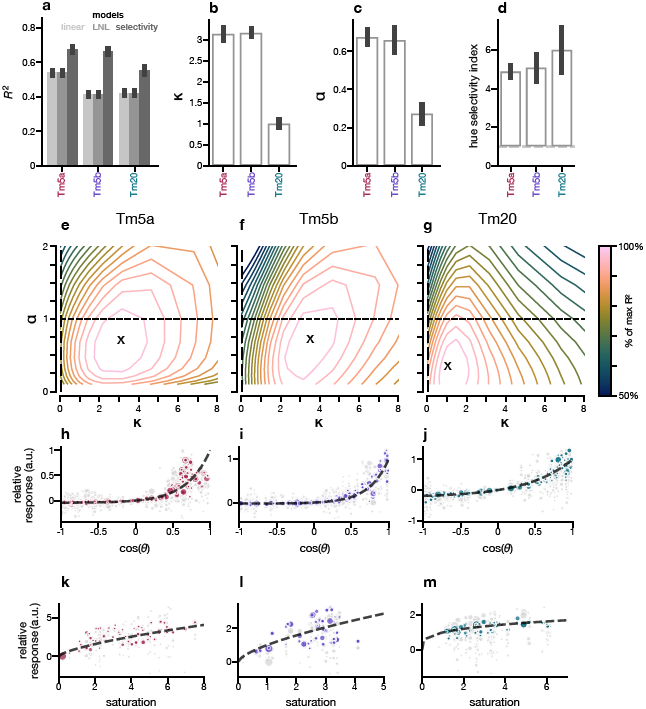
Tm5b, Tm5a and Tm20 are hue selective. **(a)** Comparison of *R*^2^ values for the linear, linear-nonlinear and the selectivity model fits to the data. The height of each bar is the *R*^2^ obtained using all the data, and the error bar is the standard deviation from fitting to bootstrap iterations of the data. **(b)** *κ* values from fitting the selectivity model. The fits are shown in (a). The height of each bar is the *κ* obtained using all the data and the error bar is the standard deviation from fitting to bootstrap iterations of the data. **(c)** *α* values from fitting the selectivity model. The height of each bar is the *α* obtained using all the data and the error bar is the standard deviation from fitting to bootstrap iterations of the data. **(d)** Hue selectivity index fitting the selectivity model. The hue index was calculated using the estimated *α* and *κ* values in (b) and (c) (see Methods). The height of each bar is the hue selectivity index obtained using all the data and the error bar is the standard deviation from fitting to bootstrap iteration of the data. **(e-g)** Effect of changing *α* and *κ* on the *R*^2^ value for Tm5a, Tm5b, and Tm20. The x marks are the mean *α* and *κ* values from panels (b) and (c). Each subsequent contour line corresponds to a reduction of the *R*^2^ relative to the maximum *R*^2^ value shown in panel (a) for the selectivity model. The dashed lines correspond to the linear regimes for *α* and *κ*. The points where the dashed lines cross represents a linear model. **(h-j)** Relative responses of the selectivity model (colored) and the data (gray) across cos(*θ*), the cosine of the angle between the preferred tuning direction and the stimulus direction. The dashed line is the fitted curve using the obtained *κ*. **(k-m)** Relative responses of the selectivity model (colored) and the data (gray) across saturation values. The dashed line is the fitted curve using the obtained *α*. Only responses with a cos(*θ*) greater than 0.5 are shown.

**Figure 4.**
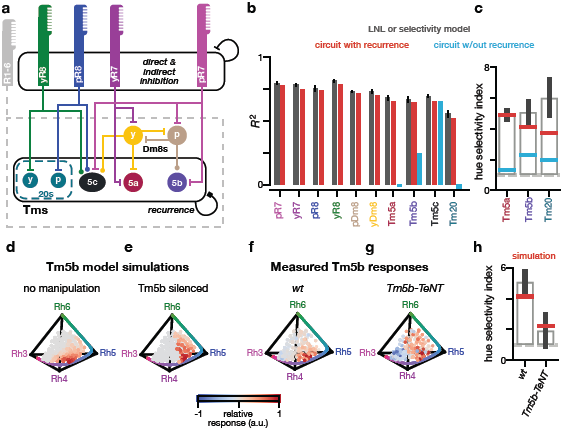
Recurrence is required for color tuning and hue selectivity of Tm5a, Tm5b, and Tm20. **(a)** Schematic of medulla color circuit indicating connections between different neurons. We have depicted two “types” of Tm20s to show connectivity specific downstream of yR8 or pR8 but our line is expressed in all Tm20s (dashed line). A connection with a flat bar is inhibitory, a connection with a filled circle is excitatory, and a connection with a filled square can be either sign. R1-6 provide indirect connections to Dm8s and Tm neurons. The connection to Dm8s is excitatory while the sign to Tm neurons is not fixed but determined by the fitting procedure. There are mono(direct) and disynaptic (indirect) recurrent connections between Tm20 and Tm5c and between Tm5a, Tm5b, and Tm5c. **(b)** *R*^2^ for the LNL or selectivity model as in figures 3a and S2a, the circuit model when fitting to all wild type data (red), and when removing recurrence in Tm neurons (cyan). **(c)** Hue selectivity index obtained by the selectivity model for Tm5b, Tm5a, and Tm20 as in figure 3d (grey bars), and the obtained selectivity index when fitting this model to the circuit model with and without recurrence (red and cyan, respectively). **(d-e)** Predicted responses of Tm5b in the full model circuit and with Tm5b silenced. **(f-g)** Measured Tm5b responses in wild type flies and in flies where TeNT is expressed in Tm5b neurons. **(h)** Predicted hue selectivity indices for Tm5b in the circuit model (simulation in red) and experimental hue selectivity indices (extracted from the data in grey) from wild type and Tm5b expressing TeNT.

This selectivity model generated fits much better than any other model for Tm5a, Tm5b and Tm20 (Fig.3a). The estimated *κ* values for Tm5a and Tm5b are around 3 and for Tm20 around 1 (Fig.3b). The estimated *α* values are slightly above 0.6 for Tm5a and Tm5b and 0.25 for Tm20 (Fig.3c). These values suggest that Tm5a and Tm5b achieve hue selectivity differently from Tm20, the former mostly through increased sensitivity to hue and the latter through decreased sensitivity to saturation.

To quantify hue selectivity, we constructed a metric equal to the ratio of the fitted model’s sensitivity to color angle (the derivative of the response with respect to cos(*θ*)) and to its sensitivity to saturation (the derivative with respect to *s*), evaluated at *θ* = 0 and *s* = 1. This metric is equal to 1 in the linear case. Our fits to the Tm5a, Tm5b and Tm20 responses all give values greater than 4, indicating nonlinearly enhanced hue selectivity (Fig.3d).

Next, we wanted to assess the sensitivity of the model to changes in the *κ* and *α* values to understand the importance of each parameter. When varying *κ* and *α*, we find that Tm5a and Tm5b are more sensitive to changes in *κ* compared to *α* whereas Tm20 is more sensitive to changes in *α* (Fig.3e-g). This further supports our hypothesis that Tm5a and Tm5b achieve hue selectivity by increasing hue sensitivity whereas Tm20 achieves hue selectivity by decreasing saturation dependence.

To visualize the tighter color tuning of the Tm neurons, it is useful to project their responses onto a twodimensional plot of response versus cos(*θ*) similar to figure 1d (Fig.3h-j). The scatter in these plots reflects both response variability and the fact that we have projected responses across a range of saturations onto this single graph. In a linear model, the responses in these plots would look linear in cos(*θ*) (Fig.1d). Instead we see responses, especially for Tm5a and Tm5b, that rise sharply near the preferred color direction (cos(*θ*) = 1). Similarly, we can visualize the tuning of Tm neurons with respect to saturation for stimuli that are close to the preferred direction of tuning (cos(*θ*) > 0.5) (Fig.1d). We find that the responses look approximately linear for the saturation plots of Tm5a and Tm5b, whereas the responses are almost flat for Tm20 (Fig.3k-m).

The results of our encoding model, combined with the reduced luminance sensitivity of Tm5a and Tm5b, identifies Tm5a, Tm5b and Tm20 neurons as hue selective.

### Connectomics reconstruction reveals a highly recurrent circuit between Tm output neurons

Our encoding model is useful to quantify hue selectivity but is not informative when it comes to determining neural circuit mechanisms that underlie these signals. To identify the circuit motifs that support hue selectivity in Tm5a, Tm5b and Tm20, we next combined connectomics constrained modeling with genetic manipulations of the circuit.

Information about the identity of the presynaptic partners to the Tms was partially determined in Takemura et al. (2015), where synaptic circuits of seven columns in the medulla were reconstructed from electron microscopy (EM). However, this dataset included many unidentified neurons, often those that span more than seven columns and did not include lobula inputs/outputs. We therefore performed independent tracing of synaptic inputs and outputs to the hue selective Tms, Tm5c, as well as the amacrine-like neurons yDm8 and pDm8, major inputs to Tm5a and Tm5b respectively (Fig.S3 and Table 1) (Zheng et al., 2018; Dorkenwald et al., 2023; Schlegel et al., 2023). Overall, our results are consistent with the seven medulla column dataset, but they offer novel insights into the connectivity of this circuit.

**Table 1.** Quantification of presynaptic sites to seed Dm8 and Tm cells. **(col.1-2)** FlyWire segment id and cell type. **(col. 3)** Total number of presynaptic connections from all presynaptic neurons. **(col. 4)** Identified presynaptic sites from cells forming >2 synapses with the seed neurons. **(col.5)** Percentage of identified presynaptic sites vs total presynpatic sites. **(col. 6)** Total number of presynaptic neurons forming >2 synapses for each seed neuron. Numbers shown inside () reflect the number of neurons identified to a cell class or a cell type. When omitted all neurons have been identified to a cell class or type.

| Flywire ID | cell type | pre sites<br>total (n) | pre sites<br>identified (n) | pre sites<br>identified (%) | neurons<br>(n) |
| --- | --- | --- | --- | --- | --- |
| 720575940638634960 | pDm8 | 284 | 217 | 76.4 | 31 |
| 720575940638424895 | yDm8 | 462 | 379 | 82 | 47 |
| 720575940639473998 | Tm5a | 227 | 152 | 67 | 21 |
| 720575940614902495 | Tm5a | 140 | 107 | 76.4 | 17 |
| 720575940627282584 | Tm5b | 328 | 225 | 68.6 | 41 |
| 720575940660846721 | Tm5b | 249 | 171 | 68.7 | 31 |
| 720575940619252603 | Tm5c | 240 | 150 | 62.5 | 30 |
| 720575940627314521 | Tm5c | 280 | 227 | 81.1 | 27 (26) |
| 720575940635252890 | Tm20 | 274 | 239 | 87.2 | 24 (22) |
| 720575940628559366 | Tm20 | 270 | 217 | 80.4 | 27 |
|  | Total | 2754 | 2084 | 75.7 | 296 (293) |

The two major inputs to Tm5a are their “home” column yR7 and yDm8 (*≈* 20% each) (Fig.S3a). An equivalent circuit exists upstream of Tm5b, with pR7 and pDm8 being the major inputs (*≈* 12% each) (Fig.S3b). Dm8s are amacrine-like interneurons that have processes that cover 10-15 ommatidial columns in layer M6 of the medulla (Gao et al., 2008; Karuppudurai et al., 2014). The two subtypes are defined by the identity of the strongest single R7 subtype input in their home column (Courgeon and Desplan, 2019; Menon et al., 2019). Taking one step back to identify inputs to Dm8s did not reveal any major neurons that might influence Tm5a and Tm5b responses (Fig.S3c, d). The main inputs to these neurons are yR7 and pR7 in their home column, although they also receive inputs from other R7s irrespective of their type. As expected from previous work, Dm8 neurons synapse onto each other (Li et al., 2021), and receive indirect R1-6 inputs (Pagni et al., 2021). The rest of the inputs is dominated by large Mt neurons that connect neighboring Dm8s. In addition to these two main inputs, amongst neurons previously assigned to a function in the medulla, Tm5a and Tm5b get indirect inputs from R1-6 (through Mi4 ≈6%). Both also get direct inputs from themselves, each other and Tm5c. Together these inputs correspond to approximately 9-10% of total inputs to Tm5a and Tm5b. Amongst the remaining inputs, the local interneuron Mi3 stands out as a common input that corresponds to approximately 8% of total inputs, as well as several large medulla tangential neurons.

Connectivity onto Tm20 is simpler (Fig.S3e and f). The major inputs to Tm20 are either a single yR8 or single pR8 (*≈* 13%), as well as LMCs (*≈* 17-30%) from the same ommatidial column. The latter should provide OFF contributions from R1-6, but Tm20 also receive ON contributions from Mi4 and Mi1 (*≈* 17%). These neurons together make up more than half of all the inputs to Tm20. Tm20s also get input from themselves (*≈* 2-5%). We did not see any obvious differences in input connectivity between Tm20s downstream of yR8 and pR8.

We also reconstructed Tm5c inputs (Fig.S3g and h). The two neurons we reconstructed received either purely y column input (yR8 and to a lesser extent yR7 and yDm8) or a mixture of yR8 and pR8, and some R7.

Together with LMCs L3 and L5, these constitute around half of all inputs to Tm5c. The Tm5c downstream of y columns got inputs from yDm8. The rest is mostly feedback from the central brain through visual projection neurons.

Because of the prevalence of self connections in the circuit, we also reconstructed outputs of the Tms down to 5 synapses (Table 2). This analysis revealed that both Mi3 and several of the large medulla tangential neurons inputs, which we named Mti_M6_M8 and Mti_M6 found in the input connectivity to Tms are also outputs of the Tms (Fig.S4). Tms are therefore connected by a recurrent network that involves both of these cell types.

**Table 2.** Quantification of postsynaptic sites to seed Dm8 and Tm cells. **(col.1-2)** FlyWire segment id and cell type. **(col. 3)** Total number of postsynaptic connections from all postsynaptic neurons. **(col. 4)** Identified postsynaptic sites from cells forming >4 synapses with the seed neurons. **(col.5)** Percentage of identified postsynaptic sites vs total postsynpatic sites. **(col. 6)** Total number of postsynaptic neurons forming >4 synapses for each seed neuron. Numbers shown inside () reflect the number of neurons identified to a cell class or a cell type.

| Flywire ID | cell type | post sites<br>total (n) | post sites<br>identified (n) | post sites<br>identified (%) | neurons<br>(n) |
| --- | --- | --- | --- | --- | --- |
| 720575940638634960 | pDm8 | 487 | 278 | 57.1 | 28 (27) |
| 720575940638424895 | yDm8 | 750 | 375 | 50 | 64 (62) |
| 720575940639473998 | Tm5a | 496 | 144 | 29 | 16 (15) |
| 720575940614902495 | Tm5a | 386 | 154 | 39.9 | 19 (17) |
| 720575940627282584 | Tm5b | 563 | 156 | 27.7 | 18 (15) |
| 720575940660846721 | Tm5b | 542 | 158 | 29.2 | 21 (18) |
| 720575940619252603 | Tm5c | 908 | 253 | 27.9 | 29 (25) |
| 720575940627314521 | Tm5c | 812 | 206 | 25.4 | 22 (19) |
| 720575940635252890 | Tm20 | 620 | 236 | 38.1 | 31 (24) |
| 720575940628559366 | Tm20 | 598 | 172 | 28.8 | 26 (22) |
|  | Total | 6162 | 2132 | 34.6 | 274 (244) |

Overall, the picture that emerges from this connectomics reconstruction is that of a feedforward circuit driven largely by R7s and R8s combined with a highly interconnected circuit, where both direct and indirect connections contribute to recurrence, mostly in the form of lateral connections, at multiple levels, a factor that was underestimated in the original seven-column dataset. A simplified version of this chromatic circuit is depicted in Fig.4a.

### yDm8 and pDm8 response are linear but different from their primary inputs

Synaptic connectivity highlights pDm8 and yDm8 as important nodes in the color processing circuit. To incorporate them into our circuit constrained model, we measured the responses of these interneurons to the same stimulus set that we used for the photoreceptors and Tms.

We found that pDm8 is excited by combinations of Rh4, Rh5 and Rh6 activation, with larger responses in the area of the tetrahedron corresponding to violet (Fig.S1e). In addition, it has inhibitory responses along the Rh3/Rh4/Rh6 face of the tetrahedron. yDm8 is broadly inhibited, along the Rh3/Rh4, Rh3/Rh6 and Rh4/Rh5 edges. It has positive responses close to the Rh6 vertex and along the Rh5/Rh6 edge (Fig.S1f). pDm8 and yDm8 responses peak along the single wavelength line (Fig.S1m-n), in agreement with previous recordings using isoquantal single wavelength stimuli (Li et al., 2021), and are not non-spectral (Fig.S1s-t). The overall direction of the tuning of Tm5b aligns with that of pDm8 (Fig.S1e). However, we find that Tm5a tuning is flipped relative to yDm8 tuning (Fig.S1f). This suggests that pDm8 forms excitatory synapses with Tm5b and inhibitory synapses with Tm5a. This is plausible because Dm8 is glutamatergic (Davis et al., 2020; Li et al., 2021) and, in fruit flies, glutamate can be either excitatory or inhibitory depending on the postsynaptic receptor. The sparsity indices for pDm8 and yDm8 are 0.65 and 0.55, respectively (Fig.S1h), and their luminance sensitivity indices are 1.25 and 0.7 (Fig.S1g). Both pDm8 and yDm8 responses are accurately predicted by a linear model (Fig.S2a). Adding an output non-linearity does not improve the fits. These results suggest that Tm5a and Tm5b may inherit some luminance invariance, but not their hue selective properties, from Dm8 interneurons.

### Recurrence is required for hue selectivity in Tm5a, Tm5b, and Tm20

With information about synaptic connectivity and measurements of the neural responses of all the main neurons in this circuit to a quasi full color space of stimuli, it is possible to now ask what features of the circuit contribute to hue selectivity. To this end, we built a circuit model constrained by connectomics data that can account for the observed responses in all the neurons we imaged - color opponent photoreceptor terminals, interneurons and hue selective Tms. This recurrent circuit model is based on known direct and indirect connections in the medulla connectome (Fig.4a). In this model, each neuron, including Tms, integrates its inputs linearly and applies a tanh output nonlinearity with a free offset parameter that determines when the neuron saturates. This is a key difference between the connectomics constrained model and the encoding selectivity model, where Tm neurons explicitly integrate photoreceptor signals non-linearly. We fit the circuit model to the responses of all neurons in three stages. In the first stage we fit the parameters of the photoreceptor-Dm9 circuit, in the second stage those for the Dm8 recurrent circuit, and the last stage the parameters of the Tm recurrent circuit.

In Heath et al. (2020), we showed that opponent responses in photoreceptor axons can be accurately modelled using EM-derived synaptic counts as quantitative estimates of the synaptic weights. Thus, we fix the weights between R7s, R8s, and Dm9s to values proportional to the connectome synapse counts, as in Heath et al. (2020), and included two gain parameters one common gain for all of the photoreceptor axons, and one for the Dm9 interneuron. A third gain parameter was added to vary the overall gain of the driving R7 and R8 rhabdomeric inputs. In total the first stage of the model has 7 parameters, three gains and four offsets. This corresponds to fitting 1.75 parameters per neuron, which is less than the four parameters per neuron used in the linear model.

The second stage of the circuit corresponds to the indirect connection between R1-6 and Dm8s, direct connections from R7s onto Dm8s, and recurrent connections between Dm8s. We fit the proportion of the R1-6 inputs and the recurrent connections between Dm8s, as well as a common gain parameter for pDm8 and yDm8. We constrained R1-6 inputs to be positive and R7 inputs and indirect connections to be negative, as shown previously (Pagni et al., 2021; Li et al., 2021). Including offset parameters, there are a total of seven parameters that we fit to the Dm8 data. This corresponds to fitting 3.5 parameters per neuron, which is less than in the four-parameter linear model.

In the final stage of the circuit corresponding to the feedforward and recurrent connections onto Tm neurons, we fixed the feedforward weight from R7s, R8s, and Dm8s as well as the recurrent weights between Tm neurons according to the relative synaptic counts of our EM reconstruction. The total weight for each recurrent connection included both direct (monosynaptic) and indirect (disynaptic) connections as identified in the EM reconstruction. We fit the proportion of the R1-6 inputs onto each Tm neuron, a separate gain for each Tm neuron, and the offset parameter for the nonlinearity. This corresponds to a total of 12 parameters or fitting 3 parameters per neuron, which is less than the 4 parameters of the linear model and the 6 parameters of the selectivity model.

The connectome-constrained circuit model fits the measured responses of R7s, R8s, Dm8s and Tm5c as well as the linear models, and Tm5a, Tm5b, and Tm20 as well as the phenomenological nonlinear selectivity model (Fig.4b) despite having fewer parameters. Across the color gamut, simulated responses from our circuit model are strikingly similar to the measured responses (Fig.S5a-j). We computed hue selective indices for Tm5a, Tm5b and Tm20 in the circuit model and obtained values comparable to those obtained directly from the data (Fig.4c).

The unexpected prevalence of recurrence in the circuit hinted at an important role for these connections to establish the hue selective responses of Tm5a, Tm5b and Tm20. We tested this hypothesis *in silico* first by removing all recurrent connections between Tms in the circuit model. In this scenario, the color tuning of Tm5a, Tm5b and Tm20 changed and, in addition, their responses became much broader (Fig.S5l-p). This result shows that recurrence between Tm neurons is critical for hue selectivity in the model. We therefore hypothesized that recurrent connections are essential to tune and sharpen the responses of these Tm neurons.

We cannot disrupt all recurrence at the level of Tms experimentally. We can however partially disrupt recurrence by blocking the output (but not the response) of a single Tm type *in vivo*. We performed this experiment by expressing Tetanus neurotoxin (TeNT) in Tm5b, which blocks neurotransmitter release (Sweeney et al., 1995), along with the equivalent manipulation *in silico*. In the circuit model, when Tm5b outputs are removed, Tm5b tuning and selectivity was affected. The tuning shifted slightly towards the Rh5/Rh6 edge and the responses were broader (Fig.4d-e). Correspondingly, the hue selectivity index dropped from 4 to 2 (Fig.4h). Experimentally, just as in the circuit model, the preferred tuning direction of Tm5b showed a slight shift towards longer wavelengths, and responses became less selective (Fig.4f-g), with the hue selectivity index dropping to 2 (Fig.4h). In fact, the neural response became effectively indistinguishable from those of a linear model under this manipulation.

In summary, our connectomics-constrained model combined with genetic manipulation of the circuit reveals that the circuit requires no nonlinear synaptic integration to achieve hue selectivity but rather the recurrence between neurons is necessary for establishing hue selective responses.

## Discussion

Color-related signals have been measured across species that use chromatic information to drive their behaviors. However, signals that are closely related to perceptual variables, such as hue or saturation, have only been reported in the cortex of primates (Hanazawa et al., 2000; Horwitz and Hass, 2012; Conway et al., 2007; Lennie et al., 1990; Komatsu et al., 1992; Zeki, 1980). Here we have identified neurons in the optic lobe of the fruit fly that have the characteristics of hue selective neurons. These neurons respond to a narrow part of the fly color space, different from their main photoreceptor input, display an increased sensitivity to hue and/or a decreased sensitivity to saturation, and luminance invariance to a degree that varies between neurons. This finding provided a unique opportunity to define neural circuit mechanisms for the emergence of hue selective signals in visual circuits in a genetically tractable organism. Using a connectomics constrained modeling approach, combined with genetic manipulations of the circuit, we show that recurrent connections, overlaid on a simple feedforward network, are critical for establishing hue selective signals, without any need for nonlinear synaptic integration. Our findings reveal the circuit basis for a transition from physical detection to sensory perception in color vision.

Our encoding models suggest that Tm5a/b and Tm20 achieve hue selectivity in different ways. Tm5a and Tm5b show an increased sensitivity to hue as well as modest saturation invariance, whereas Tm20 is invariant to changes in saturation. This suggests that different circuit mechanisms may underlie hue selectivity in these neurons. A key difference between circuits upstream of Tm5a/b and Tm20 is the amacrine cell Dm8, presynaptic to Tm5a and Tm5b and not Tm20. Our circuit model suggests that Dm8s are critical for establishing the tuning properties of Tm5a and Tm5b. Indeed pDm8 tuning starts to point towards violet, as a consequence of inhibition from yDm8 in the long wavelengths, thereby already establishing that preferred direction for Tm5b. Dm8s may also be important in increasing Tm5a and Tm5b’s sensitivity to hue.

The implementation of nonlinearities is a critical distinction between our phenomenological encoding model and our circuit constrained model. In the encoding model, Tm neurons explicitly integrate photoreceptor signals non-linearly. Conversely, the circuit constrained model employs linear synaptic integration with nonlinearity arising only from a common saturating output nonlinearity across all elements of the circuit. The results of our study reveal recurrence as a fundamental mechanism in biological circuits that enables complex nonlinear computations to be performed despite linear synaptic integration, instead of some form of nonlinear integration (Horwitz and Hass, 2012; Mehrani et al., 2020; Silver, 2010; Hubel and Wiesel, 1962).

It is not clear whether Tm20s that receive inputs from yR8 or pR8 are molecularly different cell types, but the Gal4 line we use for Tm20 labels both “types” of Tm20. Because of the differences in R8 inputs these two types of Tm20 receive, it is likely that they have different tuning, one with a preference close to the Rh5 vertex and the other closer to the Rh6 vertex. Separate measurements of the two Tm20s may also reveal responses that are more hue selective than what we measured, with one pointing towards Rh5 and the other towards Rh6. Tm5c is not hue selective and instead responds broadly in color space and is tuned to changes in luminance. Our output reconstruction, although partial (*≈* 30% of the total number of synapse), finds that the outputs with most synapses are medulla neurons. This suggests that Tm5c, which is a glutamatergic neuron (Konstantinides et al., 2018), unlike Tm5a, Tm5b and Tm20, which are predicted to be cholinergic, may act as a local interneuron in this circuit, instead of a projection neuron, perhaps establishing some of the luminance invariant properties of the hue selective Tms.

Although animals such as birds have been shown to discriminate non-spectral colors behaviorally (Stoddard et al., 2020; Daumer, 1956), non-spectral color signals have not been measured explicitly across the animal kingdom. For the fruit fly, it is unclear what the color selectivities we report here correspond to in its visual experience. Many physiological or behavioral studies of color vision in fruit flies have focused on a very narrow part of the fly color space, consisting of “visible” blue and green spectral wavelengths (Melnattur et al., 2014; Schnaitmann et al., 2013). This might explain why color behavior in fruit flies has been notoriously difficult to assess. Our measurements suggest that, moving forward, it will be necessary to expand the range of chromatic stimuli to include violet and non-spectral colors, to uncover the role of chromatic information in driving *Drosophila* behavior.

In trichromatic primates, hue selective neurons have selectivities that finely tile color space. In theory the numbers of hues that can be encoded in a tetrachromatic animal such as *Drosophila* is larger than in a trichromatic animal, with two hue angles instead of one, defining a sphere rather than a circle. However, at the Tm level at least, the possible sphere of hues is subdivided into only three hue selective signals: Violet, a non-spectral combination of UV and green, and blue/green (or up to four if we consider two Tm20s). It is possible that these signals are further combined downstream of these Tms to give rise to additional hue selective signals. Alternatively, the fly color vision system is simpler, using few and specific hue selective signals of important ethological relevance for their lifestyle, rather than for image reconstruction.

Finally, the emergence of hue signals within the chromatic circuits of flies represents an unexpected complexity at such an early stage of visual processing. Traditionally, Tm neurons have been compared to retinal ganglion cells, due to their anatomical position within their respective circuits (Sanes and Zipursky, 2010). However, our empirical findings suggest that, at least in chromatic circuits, Tms perform a function analogous to cortical neurons despite being only one to two synapses downstream of photoreceptors. Mechanistically, the extensive recurrence observed within the medulla is instrumental for producing nonlinear (higher-order) signals in a relatively superficial/small circuit. In primates, hue selectivity first appears in the primary visual cortex V1 (Horwitz and Hass, 2012; Hanazawa et al., 2000; Lennie et al., 1990). Recurrent horizontal connections within V1 are prevalent, and have been implicated in a variety of functions, including the sharpening and contrast invariance of orientation selectivity (Ferster and Miller, 2000; Sompolinsky and Shapley, 1997). The convergent evolution of algorithmic and mechanistic level computations between vertebrate and invertebrate visual circuits is well-documented (Sanes and Zipursky, 2010; Clark and Demb, 2016), and already apparent in peripheral color circuits (Fig.S7) (Heath et al., 2020). Thus, we hypothesize that the recurrent mechanism we identified may serve as a foundational basis for hue selectivity across animal brains, including primates (Fig.S7). Our findings open up new avenues for further investigation in both invertebrate and vertebrate vision.

## Acknowledgements

We are thankful to the Princeton FlyWire team and members of the Murthy and Seung labs for development and maintenance of FlyWire (supported by BRAIN Initiative grant MH117815 to Murthy and Seung). We thank the FAFB team, especially Michael Reiser and his lab members, for providing technical support and help with the EM reconstructions on CATMAID. We thank Nick Manfred, Henrique Ludwig and Miriam Flynn for their initial work on Dm8 reconstructions. We thank Aljoscha Nern and Gerry Rubin for providing the Tm5a driver. We thank Philip Schlegel for developing and sharing the NAVis python library for connectome data analysis. We thank Richard Axel for comments on the manuscript.

MPC was supported from NIH 5T32EY013933 and NIH R01EY029311. ASD and MSW were supported by NIH R01EY029311. SLH acknowledges support from NSF GRF DGE-1644869 and NIH F31EY030319. AA was supported by the Zuckerman Institute and the Pew Charitable Trusts. L.F.A was supported by NSF NeuroNex 1707398, the Gatsby Charitable Foundation GAT3708 and the Simons Collaboration for the Global Brain. RB was supported by NIH R01EY029311, the McKnight Foundation, the Grossman Charitable Trust, the Pew Charitable Trusts, the Mathers foundation and the Kavli Foundation.

## Author Contributions

RB, MPC, and ASD conceived of experiments. RB, MPC, and LFA wrote the manuscript with input from ASD. ASD acquired the imaging data, performed animal husbandry, and performed EM reconstructions. AA performed EM reconstructions. MSW did the initial characterization of yDm8, pDm8, Tm5a and Tm5b responses. MPC provided the color theoretical work, conceived the stimulus design and built the underlying software, processed and analyzed the data, and performed the modeling work with input from LFA. SLH acquired imaging data for photoreceptors.

## Methods

### Fly Genetics

w+ flies were reared on standard molasses-based medium at 25°C. Rhodopsin Gal4 drivers were used for imaging photoreceptors Rh3-Gal4 and Rh6-Gal (Cook et al., 2003) along with Rh4-Gal4 and Rh5-Gal4 (Saint-Charles et al., 2016). Dm8 cells were targeted for imaging using *OrtC2b-GAL4,DIP*γ*-GAL80/+;* Courgeon and Desplan (2019), *R24F06-GAL4, Dip*γ*-GAL80* Courgeon and Desplan (2019) and *OrtC1-3-Vp16; DIP*γ*Gal4DBD* Courgeon and Desplan (2019). Tm cells were targeted using the following divers: Tm5a: *27E03p65ADZp attP40; 94H07-ZpGdbd attP2* (Gift from A. Nern); Tm5b: *OrtC1aDBD;ET18kdVP16AD* Karuppudurai et al. (2014); Tm20: *41E03-p65ADZp;81G11Z-pGdbd* ; Tm5c *ortC1a-G4DBD, OK371-Vp16AP* Karuppudurai et al. (2014). Synaptic transmission was blocked using a UAS-TeNT construct *PUASTeTxLC*.*tntG2* Bloomington Drosophila Stock Center (BDSC): 28838.

All constructs were expressed heterozygously along with 20X-UAS-GCaMP6f, also expressed heterozygously (BDSC: 42747, 52869).

Fly genotypes used on experiments: rh3»GCaMP: *w+; Rh3-GAL4/+; 20xUAS-GCaMP6f/+;* rh4»GCaMP: *w+; Rh4-GAL4/+; 20xUAS-GCaMP6f/+;* rh5»GCaMP: *w+; Rh5-GAL4/+; 20xUAS-GCaMP6f/+;* rh6»GCaMP: *w+; 20xUAS-GCaMP6f/+; Rh6-GAL4/+;* pdm8»GCaMP: *w+; 20XUAS-GCamp6f/+; R24F06-GAL4,Dip*γ*GAL80/+;* and *w+; 20XUAS-GCamp6f/+; OrtC2bGAL4,DIP*γ*-GAL80/+;* ydm8»GCaMP: *w+; OrtC1-3Vp16/20xUAS-GCaMP6f; DIP*γ*-GAL4DBD/+;* tm5b»GCaMP: *w+; OrtC1a-GAL4DBD,20XUASGCaMP6f/CYO; VP16AD24g/TM6B;* tm5bTeNT»GCaMP: *w+; ortC1a-G4DBD,20XUASGCaMP6f/UAS-TeTxLC*.*tnt; VP16AD24g/+;* tm5a»GCaMP: *w+; 27E03AD/20xUAS-GCaMP6f; 94H07DBD/+;* tm5c»GCaMP: *w+; OrtC1aG4DBD,OK371-Vp16AP/+; 20xUAS-GCaMP6f/+;* tm20»GCaMP: *w+; R41E03-p65ADZp/20xUAS-GCaMP6f; R81G11-ZpGdbd/+;*

### Two-Photon Calcium Imaging

Recordings were performed as previously described in Heath et al. (2020). Imaging was conducted with a twophoton microscope (Bruker) controlled by PrairieView 5.4 and a mode-locked, dispersion compensated laser (Spectraphysics) tuned to 930 nm. We imaged with a 20x water-immersion objective (Olympus XLUMPLFLN, 1.0 numerical aperture). In front of the photomultiplier tube (Hamamatsu GaAsP), we mounted a band-pass filter (Semrock 514/30 nm BrightLine) to reduce bleedthrough from the visual stimulus setup. T-Series were acquired at 15-30Hz and lasted for a maximum of ten minutes with each frame at x-y imaging being 160×60 pixels (0.58µm/pixel).

All experimental animals for functional imaging were briefly anaesthetized using carbon dioxide on the day of eclosion, and imaged at ages ranging from 3-13 days. Flies were prepared for two-photon imaging based on methods previously described (Behnia et al., 2014; Heath et al., 2020). Flies were anesthetized using ice, and mounted in a custom stainless-steel/3Dprinted holder. A window was cut in the cuticle on the caudal side of the head to expose the medulla and lobula. Photoreceptors and Dm interneurons where imaged in the medulla, whereas Tm axons were imaged in the lobula. The eyes of the fly remained face down under the holder, and remained dry while viewing the visual stimuli, while the upper part of the preparation was covered with saline. The saline composition was as follows (in mM): 103 *NaCl*, 3 KCl, 5 *n −tri(hydroxymethyl) methyl −*1*Aminoethane −sulphonic acid*, 8 *trehalose*, 10 *glucose*, 26 *N aHCO*_3_, 1 *N aH*_2_*PO*_4_, 1.5 *CaCl*_2_, and 4 *MgCl*_2_, adjusted to 270mOsm. The pH of the saline was equilibrated near 7.3 when bubbled with 95% O_2_ / 5% CO_2_ and perfused continuously over the preparation at 2 *ml/min*. The imaging region of interest was limited to the region of the medulla and lobula neurons are directly activated by stimuli. Specifically, the z-depth was zeroed at the same level for each fly (the dorsal part of the lobula) and neural responses were measured from 50-110 microns for the medulla and from 50-90 microns for the lobula below that point. Responses were measured from the rostral fourth of the medulla in that plane. The dorsal third of the eye was covered with black acrylic paint to avoid the region where Rh3 and Rh4 are coexpressed in R7s (Mazzoni et al., 2008). Calcium responses were stable throughout imaging.

### Visual Stimulation

#### Hardware

We produced full-field wavelength-specific stimuli using a customized setup (Fig.1k). The setup consists of six LEDs in the UV and visible wavelength range (ThorLabs M340L4 - dUV/340nm; M365L2 - UV/360nm; M415L4 - violet/415nm; M455L3 - blue/455nm; M565L3 - lime/565nm; M617L3 - orange/615nm). A customized driver drove the five LEDs from dUV to lime. These LEDs turned on during the return period of the x-scanning mirror in the two-photon microscope (fly-back stimulation). We used the TTL signal generated by the two-photon microscope at the beginning of each line-scan of the horizontal scanning mirror (x-mirror) to trigger the LED driver. An individual T-Cube (Thorlabs LEDD1B T-Cube) drove the orange LED. Stimuli were generated using customized software written in Python. The update rate for the LED voltage values was 180Hz.

The different light sources were focused with an aspheric condenser lens (ThorLabs ACL2520U-A) and aligned using dichroic mirrors (dUV-UV dichroic - Semrock LPD01-355RU; UV-violet dichroic - Semrock FF414-Di01; violet-blue dichroic - Semrock Di02-R442; blue-lime dichroic - Semrock FF495-Di03; lime-orange dichroic - Semrock FF605-Di02). The collimated light passed through a diffuser (ThorLabs DG10-1500A) before reaching the eye of the fly, which is positioned 2cm away.

#### Intensity calibration

In order to measure the intensity of our LEDs across many voltage outputs, we used a photo-spectrometer (250-1000nm, Ocean Optics) that was coupled by an optic fiber and a cosine corrector and was controlled using our customized Python software. The photo-spectrometer was mounted on a 3D printed holder that was designed to fit on our experimental rig and approximately aligned with the fly’s point of view. For each LED, we tested a total of 20 voltage values (linearly separated) from the minimum voltage output to the maximum voltage output. For each voltage value tested, we adjusted the integration time to fit the LED intensity measured, and averaged over 20 reads to remove shot noise.

Using the spectrometer output, we calculated the absolute irradiance (*I*_*p*_(*pλ*); in *W*/*m*^2^/*nm*) across wavelengths using the following equation:

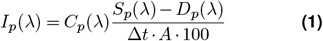

where *C*_*p*_(*λ*) is the calibration data provided by Ocean Optics (*µJ*/*count*), *S*_*p*_*(λ*) is the sample spectrum (*counts*), *D*_*p*_(*λ*) is the dark spectrum (*counts*), Δ*t* is the integration time (*s*), and A is the collection area *(cm*^*2*^). Next, we converted absolute irradiance to photon flux (*E*_*q*_; in *µE/nm*):

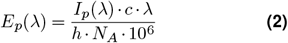

where 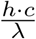 is the energy of a photon with *h* as Planck’s constant (6.63 · 10^*−*34^ *J · s*), c as the speed of light (2.998 10^8^ *m*/*s)*, and λ the wavelength (*nm*). *N*_*A*_ is Avogadro’s number (6.022 10^23^ *mol*^*−*1^).

The minimum intensity is zero for all LEDs, and the maximum intensities are (in *µE*): dUV*≈* 6, UV*≈* 7, violet*≈* 11, blue*≈* 18, lime*≈* 25, and orange*≈* 160.

#### Stimulus Design

Each stimulation protocol had at least 15 seconds before and after the stimulation period in order to measure baseline fluorescence (fluorescence to background light). Because we wanted to replicate natural spectral distributions of light, we adapted the eye to a combination of LEDs that mimic natural light conditions at dawn. We chose dawn-like conditions, because this is when flies are most active (Lazopulo et al., 2019). We fit the LED intensities to the background light condition using methods described in Christenson et al. (2022). Our background LED intensities are (in *µE*): 0.01 for dUV, 0.06 for for UV, 0.1 for violet, 0.25 for blue, 0.33 for green, and 0.25 for orange. Flies were adapted to the background light for approximately 5 minutes before the start of the recording sessions. The background light was maintained between individual recordings. During the stimulation period, individual stimuli were randomly interleaved. Each stimulus was a step stimulus that was 0.5 seconds long. The inter-stimulus interval was 1.5 or 2 seconds long.

#### Contrasts stimulus set

For the contrast stimulus set, we flashed the LEDs individually and selected mixtures of LEDs. The intensity steps were added on top of the chosen background light with added intensities of 0.1, 0.3, 0.5, 0.75, 1, and 3 µE. For LED mixtures, the intensity of each individual LED was set to these additive intensity steps, so that e.g. a mixture of violet and blue at 1µE of added intensity has a total intensity of 2µE plus the total background intensity. The LED mixtures sampled were (using the single letter annotations for the LEDs): D+U, U+L, D+U+L+O, V+B+L+O, U+V+B, B+L+O, D+L+O, D+U+V+B+L+O, V+B, D+U+O, and D+U+V. Within each imaging session we flashed all stimuli in the set once.

#### Color gamut set

For the color gamut set, we flashed different mixtures of LEDs. We obtained the LED intensities by first randomly sampling capture values in the fly color space around an overall relative capture of 5 (Fig.S6), and then fitting the target captures using methods previously described in Christenson et al. (2022). For each imaging session, we flashed a subset of this stimulus set (approx. 20% of all stimuli) and repeated each stimulus three times. We implemented a random subsampling method that ensured we span color space for each imaging session. To do this, we first randomly selected a stimulus in color space. Then, we iteratively sampled from a subset of ten not-yet-chosen stimuli that are maximally distant from the already sampled stimuli. The convex hull of the color gamut stimulus set covers approximately 90% of a set of natural reflectances within the fly’s chromatic hyperplane (Arnold et al., 2010), therefore allowing for a fairly complete characterization of a neuron’s chromatic tuning properties across the set of possible fly colors.

### Quantification of Imaging Data

All data analysis for *in vivo* calcium imaging was performed in Python using custom-made Python code and publicly available libraries. First, we removed minor remaining bleedthrough artifacts from our LED system by subtracting the 10th percentile value of each column of pixels in each image. To correct our calcium movies for motion we performed rigid translations based on template alignment using the algorithm provided by the CaImAn package (Giovannucci et al., 2019).

#### Image denoising

To denoise our calcium movies, we implemented a version of Kernel PCA that has been shown to reduce different types of noise in images (Mika et al., 1998; Weston et al., 2003). First, we reshape the movie so that all rows correspond to a frame and all columns correspond to the flattened image (*n*_*frames*_ *× m*_*pixels*_). Then, we concatenate 7 n-frame-shifted versions of the movie horizontally (*n*_*frames−*7_*×* (*m*_*pixels*_ *×*7). We perform Kernel PCA on this final data matrix using 512 components and a radial basis function. The hyperparameters of the Kernel PCA model were set to the standard values of the sklearn.decomposition.KernelPCA class function. We obtained the denoised version of the movie by taking the last m_*pixels*_ columns of the reconstructed data matrix and reshaping it back into the movie format.

#### Image Segmentation

Region of interestes (ROIs) were selected automatically using a custom-made approach and verified manually. A local correlation projection was taken of the complete motion-corrected and denoised image stack. We thresholded the projected image in three ways to identify pixels that are certainly part of a ROI (upper threshold), certainly part of the background (lower threshold), and possibly part of a ROI (medium threshold). The thresholds were chosen by fitting a two-component Gaussian Mixture model to the pixel values of the projected image using the sklearn.mixture.GaussianMixture class function. The lower and upper threshold was set to the estimated mean of the component with a lower and higher value, respectively. The medium threshold was set to the weighted average of the two means, weighted by the variance of each component. The thresholded images were used to identify connected components (i.e. individual ROIs). Next, we applied a watershed transformation to obtain the individual ROIs. We discarded any ROIs of fewer than 4 pixels.

#### Signal Extraction

To extract calcium traces from our segmented images, we first took the average fluorescence of each ROI at each time point. We subtracted the mean background fluorescence the mean fluorescence of all pixels that do not belong to any ROI from each trace to remove background fluctuations. To calculate the dF/F signal, we use as a baseline for our denoised traces the 25th percentile of a rolling 40 second time window. Finally, we smooth our dF/F signal with a Savgol filter of size 0.5 seconds and a third order polynomial. We discarded ROIs, where the signal-to-noise (SNR) ratio was smaller than 2. The SNR was defined as the magnitude of the amplitude responses during stimulation over the magnitude of the baseline responses before and after the start of stimulation (SNR = ∥*r*_*stim*_∥ ^2^/∥ *r*_*baseline*_ ∥^2^). Amplitude responses during stimulation were calculated by taking the mean dF/F signal between the 0.35 and 0.5 second during stimulation and subtracting the mean value between 0.35 and 0.1 before stimulation. Baseline responses were calculated by randomly taking mean values of a 0.15 second duration before and after the start of the stimulation protocol.

#### Response averaging and normalization

After removing noisy ROIs, we averaged the individual stimulus-aligned traces. Next, we calculated our final estimate of the amplitude using these averaged stimulus-aligned traces. As before, we take the mean dF/F signal between the 0.35 and 0.5 second during stimulation and subtract the mean value between 0.35 and 0.1 before stimulation. Next, we obtain a normalized dF/F response vector **v**^*′*^ for each neuron by dividing the unnormalized response vector (**v**) by its estimated standard deviation from zero:

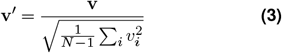

where *v*_*i*_ is the unnormalized response of a neuron to stimulus *i*, and *N* is the total number of stimuli. In all our plots where we indicate dF/F amplitudes, we show this normalized dF/F signal. For the tetrahedral plots, we show max-normalized responses to ensure equal scaling of the colored response map across neurons.

### Modeling

The stimuli we used are characterized by computed photon activations for the four fly opsins, labelled by µ = 1, 2, 3, 4, and given by *X*_*µ*_ = log ((*q*_*µ*_ + 0.001)/1.001), with *q*_*µ*_ the calculated relative capture for opsin *µ*, as described previously in Heath et al. (2020); Christenson et al. (2022). We decompose the components of the 4-dimensional vector **X** into a 3-dimensional vector 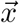 lying within the color tetrahedron and a scalar l that is the projection of **X** along the axis connecting the zero point in the full color space to the white point in the color tetrahedron. The color of each stimulus in all of our models is thus described by the 3-dimensional vector 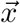, which connects the white point at the center of the color tetrahedron to the projection of the color point of the stimulus into the tetrahedron. We also define 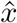 as the vector 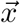 normalized to unit length. The stimuli were designed to be equal luminance but, because this could not be achieved exactly, we include a term in our models proportion to the luminance l of each stimulus. Each neuron is characterized by a three-dimensional color preference 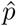, which is a vector of unit length in the color tetrahedron, an overall amplitude factor a, and the coefficient multiplying the luminance l, denoted by b. This is the full complement of parameters for the linear model, whereas there are additional parameters in the other models, as described below. Note that 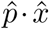 can be expressed as cos(*θ*), with *θ* the angle between 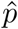 and 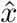, Similarly, 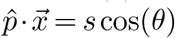, with 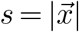 the saturation. The input for stimulus i is characterized by 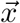 and l_*i*_, and this generates a model response y_*i*_. The corresponding response from the data is v_*i*_. All fits were done by minimizing the squared difference between y_*i*_ and v_*i*_, summed over i, except for the circuit model for which we maximize the sum of the correlation coefficients of y_*i*_ and v_*i*_ for reasons given below.

#### Linear Model

In the linear model, the predicted response to stimulus i is

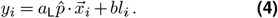

The model has 4 parameters, a_L_, b and 2 parameters that specify 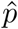.

#### Linear-nonlinear (LNL) model

For the LNL model, we first compute a linear response 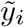 using equation 4 and then pass this through an output nonlinearity given by a modified tanh function (Rajan et al., 2009),

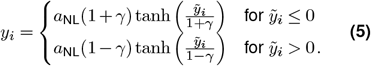

The additional parameters, beyond the 4 of the linear model, *a*_NL_ and *γ* were determined using nonlinear least-squares (`scipy.optimize.least_squares`).

#### Selectivity model

The selectivity model uses the parameters *a*_NL_, *b* and 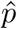 and adds two new parameters *κ* (hue sensitivity) and *α* (saturation sensitivity), with

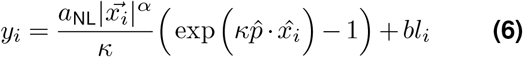

To fit this model, we optimized the parameters 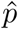, a_NL_, and b over a grid of values for *κ* and *α*, and determined the best fit across the grid. The *κ* values varied between 10^*−*2^ and 10^1^ in 20 uniform log steps, and *α* values varied between 10^*−*1^ and 10^1^ in 13 uniform log steps.

To describe the hue selectivity index, we first write equation 6 in the equivalent form

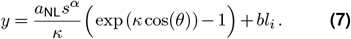

The hue selectivity index is defined as the ratio of two derivatives evaluated at s = 1 and *θ* = 0,

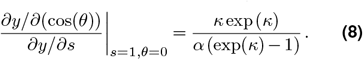

The larger this index is, the stronger the hue selectivity, and its value is 1 for the linear model.

#### Model Fitting

For all models, we used the color gamut data in our fitting procedure. Because this dataset has a different number of observations for different stimuli (see color gamut set stimulus details in Stimulus Design section), we weighted each color point during training by the number of observations. To assess goodness of fit, we calculated R^2^ values,

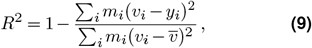

where *m*_*i*_ is the number of observations for stimulus *i*, and 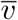 is the observation-weighted average response over all stimuli.

#### Circuit model

The circuit model is a recurrent network constrained by the existing direct and indirect connections of the EM-reconstructed medulla neuropil connectome. The prediction of the model for *a* = 1, 2,…, 11, representing pR7, yR7, pR8, yR8, Dm9, pDm8, yDm8, Tm5a, Tm5b, Tm5c, Tm20 respectively, is denoted by *y*_*a*_. These responses are given by a system of ordinary differential equations with time constants τ_*a*_,

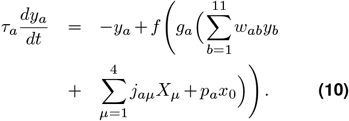

The τ values were fixed at 0.05 seconds for photoreceptors and Tm neurons, 0.025 seconds for Dm8s, and 0.0125 seconds for Dm9. The function f is the modified tanh described in equation 5 with *a*_*NL*_ = 1 and with *γ*_*a*_ a free parameter for each neuron, except for Dm9, which used the fixed value γ = 0. Thus, a total of 10 free parameters determine the *γ* values. In addition, g_*a*_ is the gain parameter for neuron *a, j*_*aµ*_ are input weights for the computed opsin activations (Heath et al., 2020), and *w*_*ab*_ is the weight of the connection from neuron b to neuron a. We also include the term *p*_*a*_*x*_0_, where x_0_ is the non-chromatic input from the R1-6 photoreceptors, and *p*_*a*_ is a neuron-specific weight for that input. We assigned equal gains to the two Dm8 neurons and for all 4 photoreceptors, so there are a total of 7 parameters characterizing the 11 neuronal gains of the model. Only 4 of the neurons have non-zero matrices *j*_*aµ*_ because opsin activations only affect photoreceptor neurons directly. The remaining 4 by 4 matrix is diagonal, and we use the same value for all 4 photoreceptors (Heath et al., 2020). Thus, the full 11 by 4 matrix *j*_*aµ*_ is characterized by a single parameter. There are only 6 nonzero values of *p* because R1-6 only synapses onto the 2 Dm8s and the 4 Tm neurons. The 11 by 11 matrix **w** (Fig.S5k) is almost entirely taken from our connectome synapse counts and from Heath et al. (2020). To obtain the weights of Tm recurrence, we added together both the mono- and di-synaptic connections between Tms. The only exceptions to a full determination of **w** are the connections between the 2 Dm8 neurons, which were not reconstructed and are thus determined by 2 additional parameters. All told, the total number of parameters is 26.

The weights from Dm8s, R7s, and R8s to Tm neurons as well as the recurrent weights between Tm neurons are fixed according to their proportional input, as obtained from our EM reconstruction. We constrained R1-6 inputs to Dm8 to be positive and R7 inputs and indirect connections to Dm8 to be negative, as shown previously (Pagni et al., 2021; Li et al., 2021). We freely varied the signs of R1-6 inputs (excitatory or inhibitory) onto the Tms, but set R7s and R8s to be inhibitory onto Tm5a, Tm5b, and Tm20 and excitatory for R8s onto Tm5c, as R8s can be both inhibitory (via histamine transmission) and excitatory (via acetylecholine transmission) (Davis et al., 2020). The signs of Dm8 inputs onto Tms were positive for Tm5b and Tm5c, and negative for yDm8 onto Tm5a. For the signs of the recurrent weights, we tested models with different signs and chose the set of signs that resulted in the best fit.

We simulated the model using discrete steps of 0.0025 seconds. We measured the simulated data as we did for our recordings. Fitting was based on the correlation coefficient *r*_*a*_ between the simulated and measured responses for each neuron. We use correlations instead of squared errors, because our calcium indicator only measured the relative amplitudes of responses. We obtain our loss function by summing the negative of the correlation over neurons,

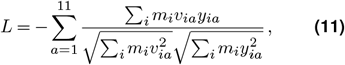

where y_*ia*_ is the response of neuron a to stimulus i. We fit the model in three stages, first fitting the parameters for the photoreceptor-Dm9 circuit, then for the Dm8 circuit, and finally the Tm circuit. For each stage, we fit the parameters using backpropagation with pytorch’s autograd functionality and its Adam optimizer. Each stage was fit over 3000 iterations with a learning rate of 0.003. Otherwise, the default parameters for the Adam optimizer were used.

#### Interpolation

To interpolate responses between sampled points in photoreceptor excitation space, we implemented a radial basis function (RBF) interpolator using a modified version of the scipy.interpolate.RBFInterpolator class. The modification was simply to remove a bias term that was not needed given that our measured amplitudes are baseline subtracted. For our RBF kernel, we chose the thin plate spline, a common spline-based technique for data interpolation and smoothing (Chui, 2001). To ensure that we were working in an interpolative regime, we projected all points outside of the convex hull of the training set onto the convex hull, using methods previously developed in the *drEye* Python package introduced in Christenson et al. (2022). We fit the interpolator by combining both the color gamut dataset and the contrast stimulus set into a single dataset. This increased the volume of the convex hull, and thus reduced clipping of interpolated responses.

For the single wavelength interpolation, we first calculated the excitations of each photoreceptor to a set of lights that follow a Gaussian distribution with a width of 10nm and peaks ranging from 320nm to 580nm. We normalized the calculated excitation for each single wavelength so that they had the same constant value as the isoluminant plane chosen for our gamut stimulus experiment. To obtain non-spectral lines, we took the set of single wavelengths and connected the single wavelength stimulus points in color space that correspond to the maximum excitation of each non-adjacent opsin.

For the single wavelength interpolation, we first calculate the excitations of each photoreceptor to a set of lights that follow a Gaussian distribution with a width of 10nm and peaks ranging from 320nm to 580nm. We normalize the calculated excitation for each single wavelength so that they have the same constant value as the isoluminant plane chosen for our gamut stimulus experiment. To obtain non-spectral lines, we took the set of single wavelengths and connected the single wavelengths stimulus points in color space that correspond to the maximum excitation of each non-adjacent opsin.

### Statistical Measures

#### Bootstrapping

Any bootstrapped distribution or calculated standard deviation shown is the result of empirical bootstrapping of independently chosen subsets of ROIs. For each bootstrap sample, we obtain mean responses as for the complete dataset and calculate the various indices and fits as before. We performed a total of 1000 bootstrap iterations to obtain our empirical distributions.

#### Sparsity index

To obtain the sparsity index, we calculate the absolute responses and normalize by the maximum absolute response and then average across these normalized responses, so that:

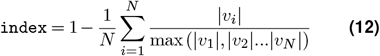

#### Luminance invariance index

To obtain the luminance invariance index, we first perform two types of regressions on the data for each neuron:

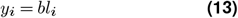

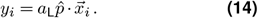

Next, we divide the obtained *R*^*2*^ value for equation 14 by the obtained *R*^*2*^ value for equation 13. We define this ratio as our luminance invariance index.

### Electron Microscopy reconstructions

We used the EM dataset from the female adult fly brain *Drosophila melanogaster* from Zheng et al. (2018) to obtain connectomic information. We chose two skeletons of Tm5a, Tm5b, Tm5c and Tm20 previously reconstructed on CATMAID (Kind et al., 2021) and found them on FlyWire (Dorkenwald et al., 2023; Schlegel et al., 2023). We accessed the FlyWire server programatically and used FAFBseg and NAvis python libraries to visualize all neurons, find all synapse locations (pre and post), list all presynaptic and postsynaptic segments connected to our seed neurons and predict the neurotransmitter identity of each cell (Buhmann et al., 2021; Eckstein et al., 2020). The connections between seed Tm neurons and photoreceptors were obtained from previously published analysis (Kind et al., 2021) and refined on CATMAID. For all other connections, after listing all the pre and postsynaptic partners to the Tm cells, we reconstructed them in the FlyWire environment, through the web interface flywire.ai, in order to identify their cell class or cell type based on their anatomy. We reconstructed and identified all presynaptic segments with >2 synapses (1488 sites) (Table 1) and postsynaptic segments >4 synapses (1479 sites) (Table 2). In the case of Dm8s, we selected “seed” pDm8 and yDm8 neurons amongst a set of previously identified cells from this volume (Kind et al., 2021) and identified them in FlyWire. We reconstructed and identified all presynaptic segments with >2 synapses (pDm8:284; yDm8:487 sites) (Table 1) and postsynaptic segments >4 synapses (pDm8:487; yDm8:750 sites) (Table 2). Home column photoreceptor presynaptic inputs had previously been identified (Kind et al., 2021), photoreceptor inputs to other columns were identified on CATMAID. Altogether, we found that our seed Tm and Dm8 neurons had 2754 postsynaptic sites and 6162 presynaptic sites. 2084 sites (75.67%) represent inputs from 296 presynaptic neurons (>2 synapses) and 2132 sites (34.6%) are outputs to 270 postsynaptic neurons (>4 synapses) (Table 2).

## Supplementary Information

**Figure S1.**
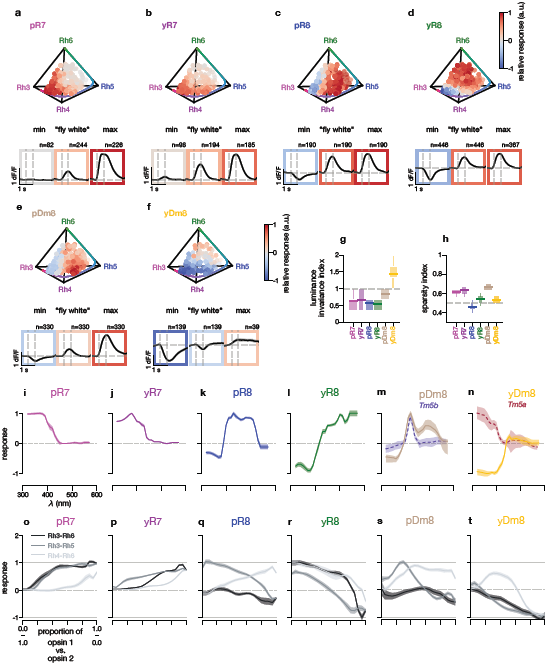
Chromatic tuning properties of R7s, R8s, and Dm8s. **(a-f)** Relative amplitude responses of R7s, R8s, and Dm8s neurons across the gamut of testable fly colors. An individual chromatic stimulus is a point in the chromatic hyperplane. The color of the point indicate the relative response of the neuron to this stimulus. The colored line that spans the edges of the tetrahedron from Rh3 to Rh6 corresponds to the optimal single wavelength line. The spectrum spans all single wavelengths from 300 to 540nm (i.e. the visible spectrum of the fly). The three insets for each plot show dF/F responses across time from 0.5s before stimulus onset to 1.5s after stimulus offset onset and offset are indicated by the dashed gray vertical lines. The dashed gray horizontal line indicates the baseline dF/F signal. From left to right, the first inset shows the average responses across recorded neurons around the location of the minimum amplitude. The second inset shows the average response around the middle of the tetrahedron the “fly white” point. The third inset shows the average response around the location of the maximum amplitude. The spherical volume used for averaging responses around the desired location for each inset has a radius of 0.05 each edge in the tetrahedron has a length of 1. **(g)** The sparsity indices for R7s, R8s, and Dm8s neurons. A value of 0.5 corresponds to a uniform response distribution. The vertical histogram corresponds to the bootstrapped distribution and the line for each cell type corresponds to the mean sparsity index. **(h)** Calculated chromatic variance indices for R7s, R8s, and Dm8s. The chromatic variance is calculated by taking the ratio of the goodness-of-fit (*r*^2^) of a linear regression using only the chromatic dimensions (i.e. only opponent-like interactions) and the *r*^2^ of a linear regression using only the achromatic dimension (i.e. sum of photoreceptors). The vertical histogram corresponds to the bootstrapped distribution and the line for each cell type corresponds to the mean achromatic variance index. **(i-n)** Interpolated single wavelengths for R7s, R8s, and Dm8s. The colored line corresponds to the interpolated mean response. The shaded areas correspond to the bootstrapped 95% percentile of the mean response. Tm5b and Tm5a single wavelength responses from Fig.2f and g, the dashed lines in m and n, are overlaid on pDm8 and yDm8 for comparison, respectively. **(o-t)** Interpolated non-spectral responses of R7s, R8s, and Dm8s. The colored lines correspond to the interpolated mean responses for different non-spectral lines (as indicated in the legend). The shaded areas correspond to the bootstrapped 95% percentile of the mean response.

**Figure S2.**
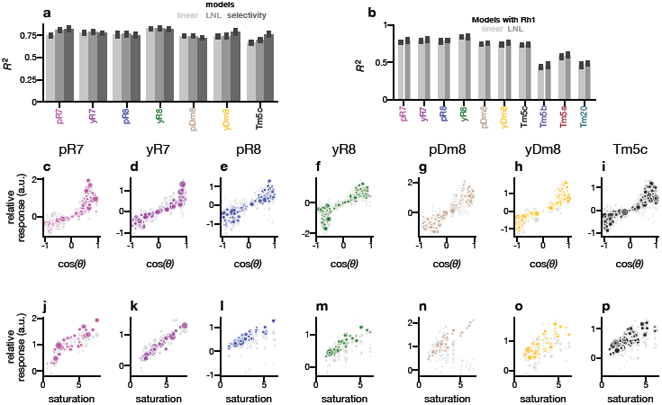
Encoding models fit to R7s, R8s, Dm8s, and Tm5c. **(a)** Comparison of different *R*^2^ values using standard linear regression, the linear-nonlinear model, and the selectivity model. The height of each bar is the *R*^2^ obtained using all the data and the error bar is the standard deviation of fitting the models to each bootstrap iteration of the data. **(b)** Obtained *R*^2^ values using a linear model with the additional Rh1 feature (i.e. the calculated excitation of R1-6 photoreceptors) for all neurons. **(c-i)** Relative responses of the LNL model (colored) and the data (gray) across cos(*θ*) - the cosine of the angle between the preferred tuning direction and the stimulus direction. **(j-p)** Relative responses of the selectivity model (colored) and the data (gray) across saturation values.

**Figure S3.**
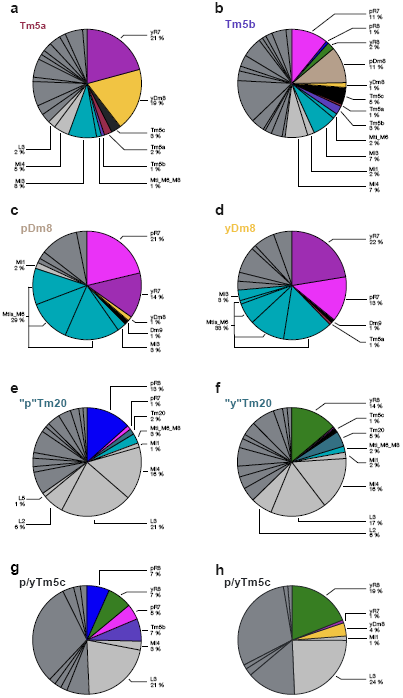
Presynaptic connectivity of Tm5a, Tm5b, Tm5c, Tm20 and y/pDm8. **(a-h)** Fraction of total inputs (>2 synapses) to Tm5a, Tm5b, Tm5c, Tm20, pDm8 and yDm8 obtained from EM reconstruction and identification of presynaptic partners in the medulla and the lobula. **(a**,**b)** Averaged fraction of inputs of two reconstructed Tm5a and two Tm5b cells respectively. **(c-h)** Input fractions onto individual neurons. Colored wedges represent inputs from neurons addressed in this study (R7s, R8s, Dm8s, Tm5s and Tm20s). Teal specifically denotes recurrent inputs (Mi3 and Mtis). Light gray wedges correspond to indirect input from R1-6 photoreceptors and dark grey are all other inputs.

**Figure S4.**
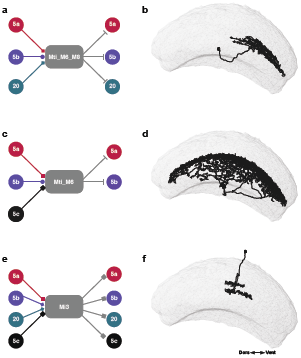
Putative chromatic recurrent circuits in the medulla between Tm cells and local interneurons. Our EM reconstructions revealed the presence of three types of interneurons that establish synaptic recurrency between different Tms. **(a**,**b)** A medulla tangential neuron was found as both pre and postsynaptic to Tm5a and Tm20. We named it Mti_M6_M8 as it ramifies across layers M6 and M8 of the medulla. Loose reconstruction of inputs and outputs from five of these cells confirmed Tm5a and Tm20 connectivity and showed that this cell type is also pre and post synaptic to Tm5b cells. Somas are located in the medial-anterior side of the medulla. Neurotransmitter prediction suggests they are GABAergic. **(c**,**d)** Tm5a, Tm5b and Tm5c send outputs to a large tangential neuron, putatively GABAergic, that expand its dendritic tree across medulla M6 layer and sends outputs to Tm5a and Tm5b, that we names Mti_M6. **(e**,**f)** Combining our Tm reconstructions and the seven column EM dataset (Takemura et al. (2015) we observed that Mi3 is both pre and postsynaptic to Tm5a, Tm5b, Tm5c and Tm20. **(b**,**d**,**f)** Single cell reconstructed skeletons of Mti_M6_M8, Mti_M6 and Mi3 (Flywire segments ids 720575940644160968, 720575940625550823, 720575940623904136 respectively).

**Figure S5.**
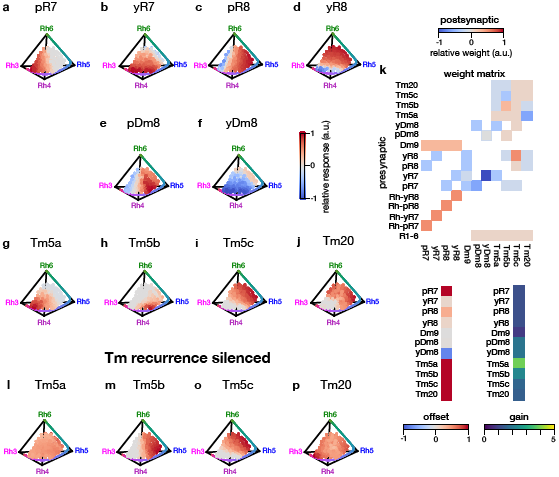
Predicted responses and weight matrix of the fitted circuit model. **(a-j)** Predicted responses of all neurons in the connectomics constrained circuit model. **(k)** Weight matrix, offset and gain parameters for the circuit model. The Rh-xRx signature indicates the calculated photoreceptor excitations of R7s and R8s at the level of the rhabdomere. pR7, pR8, yR7, and yR8 indicate the axonal segment of the photoreceptors, which we model as a separate node from the rhabdomere. All other neurons are modelled as a single node. **(i-p)** Predicted responses of Tms when silencing Tm recurrence in the circuit model.

**Figure S6.**
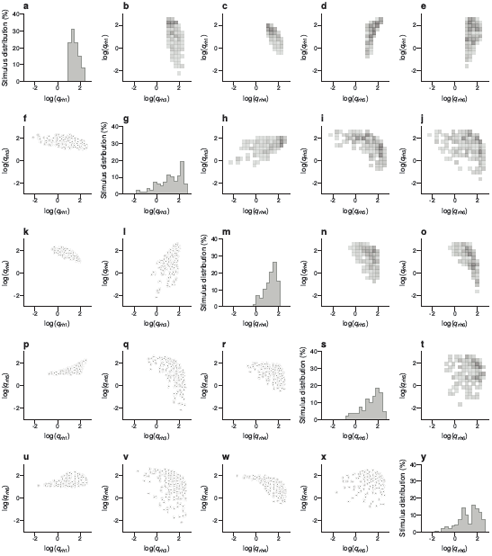
Distribution of relative capture values in the color gamut stimulus set. **(a-y)** Distribution of calculated relative captures for the Rh1, Rh3, Rh4, Rh5, Rh6 opsin. The diagonal panels show the histogram for each opsin as a percentage of the stimulus set. The panels in the lower triangle are scatter plots of the individual stimuli for pairwise comparisons between opsins. The panels in the upper triangle are 2D-histograms of paired opsins.

**Figure S7.**
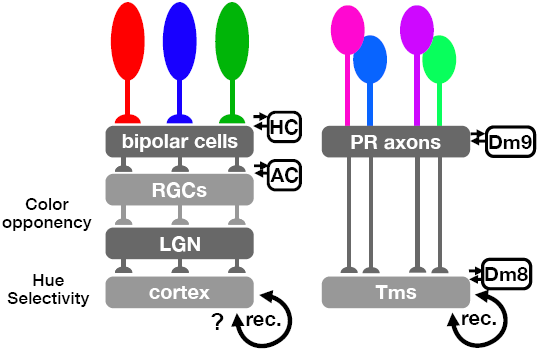
Convergent evolution of primate and fly chromatic circuits. In trichomatic primates, S, M and L cone excitation is transformed into opponent channels through feedforward and lateral processing (HC: horizontal cells, AC: amacrine cells) in the retina. The two axes of opponency - “red-green” and “blue-yellow” - have been shown to correspond to an optimal decomposition of S, M and L cone sensitivities (Buchsbaum and Gottschalk, 1983). Linear color opponent signals measured in retinal ganglion cells (RGCs) and lateral geniculate nucleus (LGN) (De Valois et al., 1966; Derrington et al., 1984) are further processed into nonlinear hue signals in cortex. Hue selective neurons have been described in V1 (Horwitz and Hass, 2012; Hanazawa et al., 2000; Lennie et al., 1990) and V4/IT (Conway et al., 2007; Komatsu et al., 1992; Zeki, 1980). In *Drosophila*, p/yR7 and R8 excitation is transformed into color opponent channels in photoreceptor axons through feedforward and lateral processing (horizontal-like cell Dm9) as well as axo-axonal interactions (Schnaitmann et al., 2018; Heath et al., 2020). The axes of opponency have also been shown to optimally decompose photoreceptor signals (Heath et al., 2020). Linear color opponent signals measured in photoreceptor axons are further processed in Dm8 amacrine-like neurons and Tm neurons where we report hue selective signals that are established through recurrent/lateral connections. Since such recurrent/lateral connections are a prevalent in the primate cortex, we propose that similar, convergent, mechanisms may also support hue selectivity in the primate cortex.

